# Schwann cells modified to secrete MANF is a potential cellular therapy for peripheral nerve regeneration

**DOI:** 10.1101/2025.03.14.642904

**Authors:** Bhadrapriya Sivakumar, Caleb Hammond, Valeria Martinez, Nickson Joseph, Johnson V. John, Anil Kumar, Anand Krishnan

## Abstract

Despite several decades of research, an effective therapy for peripheral nerve regeneration is still lacking. The lack of knowledge of molecular candidates that equally promote axon regeneration and glial cell dynamics essential for regeneration poses challenges in developing effective therapies. Improper optimization of potential therapies leading to failures in ensuring their local availability in nerves also poses additional challenges. Here, we showed that the neurotrophic factor, the mesencephalic astrocyte-derived neurotrophic factor (MANF), equally promotes axon regeneration and glial cell dynamics favorable for nerve regeneration. We showed that while endogenous expression of MANF is primarily restricted to non-peptidergic sensory neurons in adult rats, exogenous MANF promotes the growth of all subtypes of adult rat sensory neurons. We also demonstrated that exogenous MANF promotes the proliferation and migration of adult rat primary Schwann Cells (SCs). Further, we found that local and repeated administration of exogenous MANF to injured mouse nerve promote axon regeneration. Finally, we devised a therapeutic approach by programming nerve resident SCs to locally and continuously deliver MANF to injured rat nerves and showed that this approach improved nerve regeneration indices. Overall, this work developed a therapeutic approach by harnessing the power of SCs as a local delivery system of MANF for improving nerve regeneration.

## Introduction

Peripheral nerve injuries account for a significant clinical and socioeconomic burden(1). However, no available therapies effectively repair nerves and restore full functions. The lack of effective treatments often leaves the affected patients with permanent loss of function. A primary hurdle in translating experimental nerve repair therapies into effective clinical therapies is the failure to maintain their optimal local availability in regenerating nerves(2). The lack of knowledge of potent molecular candidates that can facilitate both axon regeneration and glial cell dynamics essential for peripheral nerve regeneration also challenges therapy development.

Peripheral nerve regeneration involves complex sequential steps initiated by the clearing of axonal and myelin debris from injured nerves by macrophages and Schwann Cells (SCs), the major glial cell population in peripheral nerves(3). The SCs then de-differentiate, proliferate, and migrate in an orderly manner toward the injured proximal nerve stump to form endoneurial tubes and direct regenerating proximal axons toward their target tissues(3). Clearing of debris is the most efficient process among these complex events. Although axon regeneration occurs initially and may last for several weeks to months, it gradually diminishes because of the lack of local trophic support. SCs also gradually lose their dynamics because of chronic denervation and lack of support from axons(4). Hence, even after surgical correction, nerve regeneration often fails because of the diminished growth responses from axons and SCs. Therapies that improve and maintain both axon regeneration and SC dynamics, when available locally to injured nerves, should improve regenerative outcomes.

Recent work from our lab and other researchers demonstrated that the neurotrophic factor mesencephalic astrocyte-derived neurotrophic factor (MANF) is a potential molecular candidate for improving nerve regeneration(5, 6). However, the endogenous availability of MANF in injured nerves has not been systematically explored. Similarly, whether MANF is equally effective in promoting axon regeneration and SC dynamics, which is critical for complete nerve regeneration, has not been systematically investigated. The effect of MANF in improving nerve crush injuries, the most common form of nerve injury presented in neuro-clinics, has also not been examined. In this work, we demonstrate that MANF is preferentially expressed in a subtype of sensory neurons known as the small calibre non-peptidergic sensory neuron subtype, while exogenous MANF improves the outgrowth of all subtypes of adult sensory neurons regardless of their endogenous MANF expression status. We found that MANF is sparsely expressed in SCs. We further demonstrated that direct and repeated delivery of MANF to injured nerves promote axon regeneration. Finally, we developed a potential therapeutic approach consisting of programmed SCs to locally deliver MANF to injured nerves. We demonstrated that this approach effectively improves the indices of nerve regeneration and worth following up for therapy development.

## Materials and Methods

### Sciatic nerve injury model

All animal experiments were approved by the University of Saskatchewan animal research ethics board. Four to six-week-old adult male SD rats were used for the sciatic nerve transection experiments, and 4–6-week-old adult male CD1 mice were used for *in vivo* sciatic nerve crush experiments. The animals were purchased from Charles River Laboratories, Canada. For nerve transection experiments, the sciatic nerve of rats was exposed, and a blunt transection made using fine scissors. Post operative analgesia was provided by administering sustained-release (SR) buprenorphine. Three days after the injury, the lumbar (L4, L5, and L6) DRGs were harvested from the ipsilateral side for individual neuron cultures, immunostaining, and western blot experiments. The proximal and distal nerve segments of the transected nerves were also harvested after three days of injury for western blot and immunostaining experiments.

For the nerve crush injury model, the sciatic nerve was exposed, and an injury was made by crushing the nerve using fine forceps for 15 seconds. MANF (2 µg/25µl PBS) was applied immediately to the crushed site. The muscles and skin of the animals were sutured back, and the animals were maintained under analgesia with SR buprenorphine. On day 3 and day 5, the sciatic nerve was re-exposed for MANF treatment. Saline was applied to the control group. On day 7, the animals were euthanized. The sciatic nerves were harvested, fixed and immunostained for βIII tubulin for quantification of regenerating axons using ImageJ.

### Immunohistochemistry

The DRGs and sciatic nerves were fixed in Zamboni’s buffer (2% paraformaldehyde (Cat No. F79-500, Fisher Scientific) and 0.5% picric acid (Cat No.5860-16, Ricca Chemical) in PBS) overnight at 4^°^C, followed by incubation in 20% sucrose (Cat No. BP220-1, Fisher Scientific) for 24 h at 4^°^C. Tissue blocks were prepared using an optimal cutting temperature compound (Cat No. 23-730-571, Fisher Scientific) and 12 µm sections were collected onto slides using a cryostat. The sections were stored at −80^°^C until further analysis. For immunostaining, the sections were blocked using an immunoblocker (5% donkey serum (Cat No. 56-646-05ML, Fisher Scientific) and 0.3% Triton-X (Cat No. T8787-100ML, Sigma Aldrich) in PBS) for 40 minutes, washed in PBS and then incubated with primary antibodies for 1h at RT. The primary antibodies used were MANF (Cat No. SAB3500384, Millipore Sigma, 1:100 dilution), NF200 (Cat No. N0142, Millipore Sigma, 1:100 dilution), βIII tubulin (Cat No. MAB1637, Millipore Sigma, 1:100 dilution), GFAP (Cat No. PA1-10004, ThermoFisher Scientific, 1:100 dilution), and GAP43 (Cat No. PA5-143568, ThermoFisher Scientific, 1:100 dilution). The secondary antibodies used were goat anti-rabbit Alexa Fluor 546 (Cat No. A-11035, ThermoFisher Scientific, 1:100 dilution), goat anti-mouse Alexa Fluor 647 (Cat No. A-21235, ThermoFisher Scientific, 1:100 dilution), goat anti-rabbit Alexa Fluor 488 (Cat No. A-11034, ThermoFisher Scientific, 1:100 dilution), and goat anti-chicken Alexa Fluor 647 (Cat No. A-21449, ThermoFisher Scientific, 1:100 dilution). The secondary antibodies were incubated for 1h at RT. For IB4 positive neuron staining, the sections were initially incubated with Griffonia Simplicifolia Lectin 1 (GSL1) Isolectin B4 (IB4) (Cat No. L-1104-1, Vector Laboratories, 1:100) for 1h at RT followed by incubation with goat anti-Griffonia Simplicifolia Lectin I antibody for 1h at RT (Cat No. AS-2104-1, Vector Laboratories, 1:100 dilution). The sections were then developed using rabbit anti-goat Alexa Fluor 488 (Cat No, A- 11078, ThermoFisher Scientific, 1:100 dilution). The sections were mounted using SlowFade Diamond Antifade mountant with DAPI (Cat No. S36973, ThermoFisher Scientific). The images were taken using a Zeiss Axio Observer inverted fluorescence microscope.

### Western Blot

Proteins were isolated using RIPA buffer (Cat No. PI8990, Fisher Scientific) in the presence of a halt protease and phosphatase inhibitor cocktail (Cat No. PI78441, MilliporeSigma). The protein concentration was determined using DC^TM^ protein assay kit II (Cat No. 5000112, Bio-Rad) according to the manufacturer’s instructions. 20μg of total protein was used for SDS-PAGE and the resolved proteins were transferred onto a PVDF membrane (Cat No. 1620177, Bio-Rad) using a semi-dry transfer method. The membrane was then blocked for 1h in blocking buffer (5% skim milk powder and 0.1% Tween-20 (Cat No. BP337-500, Fisher Scientific) in TBS). The primary antibodies used were MANF (Cat No. SAB3500384, Millipore Sigma, 1:1000 dilution) and GAPDH (Cat No. SAB3500247, Millipore Sigma, 1:1000 dilution) for overnight incubation at 4^°^C. The secondary antibodies used were goat anti-rabbit HRP (Cat No.1706515, Bio-Rad, 1:3000 dilution) and goat anti-chicken HRP (Cat No. A16054, ThermoFisher Scientific, 1:3000 dilution) for 1h at RT. The blots were developed using an ECL substrate (Cat No. 1705060, Bio-Rad) and imaged using a GelDoc (Bio-Rad).

### Neurite outgrowth analysis

Primary sensory neurons were cultured using injured and normal DRGs, as described previously(7, 8). Briefly, individual cells were enzymatically dissociated from DRGs by incubating the DRGs in 0.1% collagenase (Cat No. 17104019, ThermoFisher Scientific) for 90 minutes at 37^°^C. This was followed by the mechanical dissociation of cells from the DRGs by repeated manual pipetting. The resulting cell suspension was centrifuged at 800 rpm for 6 minutes. The cell pellet was suspended in L15 media and then layered over 15% bovine serum albumin (Cat No. SH3057402, Fisher Scientific) and centrifuged at 800 rpm for 6 minutes. The middle layer of debris in the resulting supernatant was carefully removed, and the pellet was washed using L15 media. Finally, the pellet was resuspended in primary neuron culture media comprising DMEM/F12 (Cat No. 11- 330-057, ThermoFisher Scientific), N2 Supplement (Cat No. 17502048, ThermoFisher Scientific, 1:100 dilution), 0.1% BSA (Cat No. SH3057402, ThermoFisher Scientific), 100 ng/ml NGF (Cat No. 13257-019, ThermoFisher Scientific) and antibiotic antimycotic solution (Cat No. SV3007901, Fisher Scientific, 1:100 dilution). An equal number of neurons were seeded into Nunc LabTek II 4-well Chamber Slide (Cat No. 154453, ThermoFisher Scientific) coated with 0.01% poly-l-lysine (Cat No. A005C, Fisher Scientific) and 10 μg/ml laminin (Cat No. 23017015, ThermoFisher Scientific). To study the growth-promoting effect of MANF, 50ng/ml or 100ng/ml recombinant human MANF (Cat No. 3748-MN-050, R&D Systems) was supplemented to neuron cultures. PBS was used as the control treatment. After 24 and 48 h, the cells were fixed using 4% paraformaldehyde (Cat No. P0018500G, ThermoFisher Scientific), blocked using immunoblocker (5% donkey serum (Cat No. 56-646-05ML, Fisher Scientific) and 0.3% Triton-X (Cat No. T8787- 100ML, Sigma Aldrich) in PBS) and immunostained for βIII tubulin (mouse monoclonal, Cat No. MAB1637, Millipore Sigma, 1:100 dilution) and NF200 (mouse monoclonal, Catalogue No. N0142, Millipore Sigma, 1:100 dilution). Goat anti-mouse Alexa Fluor 488 (Cat No. A-11001, ThermoFisher Scientific, 1:100 dilution) was used as the secondary antibody. Neurite images were captured using a Zeiss Axio Observer inverted fluorescent microscope. Quantification of neurite outgrowth was done using WIS-NeuroMath Software(9).

### Primary SC cand S16 culture

Sciatic nerves were collected from adult male SD rats. The epineurium was stripped off using fine forceps, and the nerve teased out into individual fibres. The fibres were then triturated, suspended in in 5 ml DMEM-D-Valine (Custom media, Boca Scientific) and centrifuged at 800 rpm for 5 minutes at 4^°^C followed by incubation of the pellet in 1mg/ml collagenase (Cat No. 17104019, ThermoFisher Scientific) for 60 minutes at 37^°^C. The fibres were then triturated using a flame-polished Pasteur pipette and incubated in 2.5% trypsin (Cat No. T4799-5G, Sigma-Aldrich) at 37^°^C for 30 minutes. The fibres were again triturated using a flame-polished glass pipette and passed through a 40 μm cell strainer (Cat No. 22-363-549, Fisher Scientific) into a falcon tube containing 2 ml of FBS. 25ml of DMEM-D-valine was added to the filtrate and centrifuged for 5 minutes at 800 rpm at 4^°^C. The pellet was then resuspended in SC media and seeded into culture flasks coated with 0.01% poly-l-lysine (Cat No. A005C, Fisher Scientific) and 10 μg/ml laminin (Cat No. 23017015, ThermoFisher Scientific). The SC media was comprised of DMEM-D-Valine (Custom media, Boca Scientific), 10% FBS (Cat No.12483020, ThermoFisher Scientific), antibiotic antimycotic solution (Cat No. SV3007901, Fisher Scientific, 1:100 dilution), 2 mM glutamine (Cat No. 25030081, ThermoFisher Scientific), N2 supplement (Cat No. 17502048, ThermoFisher Scientific, 1:100 dilution), 10 μg/ml bovine pituitary extract (Cat No. 13028014, ThermoFisher Scientific), 5 mM forskolin (Cat No. F3917, Sigma-Aldrich) and 50 ng/ml heregulin (Cat No. 10003100 UG, Fisher Scientific).

The SC line S16 was purchased from the ATCC (USA). These cells were maintained in DMEM (Cat No. 30-2002, ATCC) containing 10% FBS (Cat No.12483020, ThermoFisher Scientific) and antibiotic antimycotic solution (Cat No. SV3007901, Fisher Scientific, 1:100 dilution). The cultures were maintained in a CO_2_ incubator at 37^0^C.

### Cell proliferation assay

SC proliferation was evaluated using MTT (3-(4,5 dimethylthiazol-2-yl)-2,5-Diphenyltetrazolium bromide) assay. Briefly, 5×10^3^ primary SCs or S16 cells were seeded into a 96-well plate previously coated with 0.01% poly-l-lysine and 10 μg/ml laminin. MANF was supplemented to these cells after 24 h. PBS was used as the control. After 24 h, the media was replaced with fresh media containing 500 μg/ml MTT (Cat No. M6494, ThermoFisher Scientific). After 4 h, the generated purple formazan crystals were dissolved in dimethyl sulfoxide (Cat No. BP231-1, Fisher Scientific). The absorbances were read at 570nm (test) and 630nm (background) using a SpectraMax M2 plate reader.

### Scratch Assay

Scratch Assay was performed to evaluate the migration of SCs. 5×10^4^ primary SCs were seeded into cell culture plates previously coated with 0.01% poly-l-lysine and 10 μg/ml laminin. When the cells reached 90% confluency, a longitudinal scratch was made in the mid area of the plate using a p1000 pipette tip and the media was replaced with fresh media containing recombinant MANF. PBS was used as the control. Brightfield images of the scratch were taken at 0, 24 and 48h time points using the AxioObserver inverted microscope. The areas of the scratches were measured using ImageJ software to evaluate the scratch closure and migration efficiency of SCs.

### Transwell Migration Assay

A transwell migration assay was performed to evaluate the migration of SCs in response to the tropic signals from MANF. Briefly, 5×10^4^ primary SCs or S16 cells were seeded into cellQART 24-well culture insert membranes (Cat No. 9318012, Sterlitech Corporation). The carrier plate was filled with media containing 50 ng/ml MANF. After 48 h, the top surface of the membrane was scraped using a sterile cotton swab to remove the cells on the top surface. The insert was then washed with PBS. The membrane was then carefully cut out using a scalpel, inverted and mounted onto a Superfrost Microscopic Slide (Cat No. 12-550-15, Fisher Scientific) using DAPI-containing mounting media. Images of DAPI-positive cells were taken using the AxioObserver inverted microscope. The number of cells that migrated on to the lower surface was counted using ImageJ.

### Preparation of lentivirus carrying Doxycycline (Dox)-inducible MANF construct

A codon optimized gene construct for human MANF was synthesized and cloned into pUC57 plasmid between EcoRI and NotI restriction sites by GenScript. The gene was excised from the pUC57 plasmid with EcoRI and NotI restriction enzymes and cloned into a Dox-inducible lentivirus plasmid pLVX TetOne Puro (Takara Bio) to generate the pLVX TetOne Puro MANF construct. The sequence of the plasmid was verified by nanopore-based sequencing with Plasmidsaurus. Lentiviruses carrying the MANF gene was produced by transducing HEK 293FT cells with pLVX TetOne Puro MANF plasmid along with the packaging plasmids pMD2.G and psPAX2 using polyethylenimine (PEI). Two days later, the cell culture supernatant containing the lentiviruses were harvested and stored at −80°C until use.

### Generation of SCs expressing Dox-inducible MANF

2 x 10^5^ primary SCs were transduced with 100μl of either lentivirus generated with pLVX TetOne Puro (lenti-empty vector) or pLVX TetOne Puro MANF (lenti-MANF) in the presence of 8 μg/ml polybrene (Cat No. TR-1003-G, Millipore Sigma). The cells were treated with 1 μg/ml puromycin (Cat No. P8833-25MG, Millipore Sigma) after 24 h to select stably transduced positive cell clones. Dox-inducible expression of MANF was evaluated in 80% confluent stably transduced cells after supplementing them with 1 μg/ml Dox (Cat No. AAJ6042203, ThermoFisher Scientific) for 24 h. The media was collected for ELISA assay to measure MANF secretion.

### ELISA

ELISA was performed using the commercially available MANF ELISA Kit (Cat No. EKL58888- 96T, Biomatik) according to the manufacturer’s instructions. Briefly, the standards and test samples were incubated in the wells for 90 minutes at 37^0^C. The solutions were then removed and 100 μl detection reagent A was added to the wells and incubated for 45 minutes at 37^0^C. Followed by the wells were washed with wash buffer for five times. After this step, 100μl detection reagent B was added to the wells and incubated for 45 minutes at 37^0^C. The wash step was repeated and 90 μl of TMB substrate was added and incubated for 20 minutes at 37^0^C. Finally, a 50μl stop solution was added to the wells, and the absorbance was read at 450 nm immediately.

### DRG-nerve explant crush injury model

DRGs that were intactly connected with a 3-4 mm peripheral nerve were harvested from adult male SD rats. A crush injury was made to the nerve *in vitro*, as described above, ∼2 mm distally to the DRG. The crushed DRG-nerves were then incubated with either lenti-empty vector or lenti-MANF for 4h in a CO_2_ incubator. Thereafter, the DRG-nerve preparations were embedded in Cultrex basement membrane extract (Cat No. 3433-010-01, R&D Systems) in a 24-well plate and supplemented with Dox (1µg/ml). The tissue explants were harvested on day 6 and day 15. The 6- day timepoint explants were immunostained for βIII tubulin *(captures protected and regenerating axons)* and MANF, while the 15-day timepoint explants were immunostained for GAP43 *(captures regenerating axons)*, GFAP, and MANF. The number of axons in the proximal *(from DRG to the crush site)* and distal *(from crush site to the distal end)* parts of explants was counted using ImageJ.

## Results

### MANF is upregulated in injured DRGs and preferentially expressed in non-peptidergic sensory neurons

Our recent proteomics work showed that MANF is upregulated in *in vitro* and *in vivo* growth primed DRGs (5). To further validate this observation, we examined the expression of MANF in non-injured and injured adult DRGs *(ipsilateral DRGs from adult SD rats after 3 days of sciatic nerve transection)* using western blot and found a clear trend *(not statistically significant)* in its upregulation in injured DRGs (**Figure 1A, B**). DRGs are primarily composed of multiple subtypes of sensory neurons and satellite glial cells, and we earlier reported that MANF is preferentially expressed in a subtype of DRG sensory neurons that poorly express NF200 (NF200^poor^ neurons) (5). Here, we further characterized MANF-expressing neurons in the DRGs. NF200 high expression marks large calibre neurons that project myelinating axons, whereas Isolectin B4 (IB4) staining marks small calibre non-peptidergic neurons that project nociceptors. Co-immunostaining of NF200, IB4 and MANF in non-injured and injured DRGs showed that MANF is preferentially expressed in the IB4^+^ neuronal subtype (**Figure 1C**). Thus, it is likely that the moderate upregulation of MANF observed in injured DRGs is mostly a reflection of its upregulation in injured IB4^+^ neurons.

**Figure 1:**
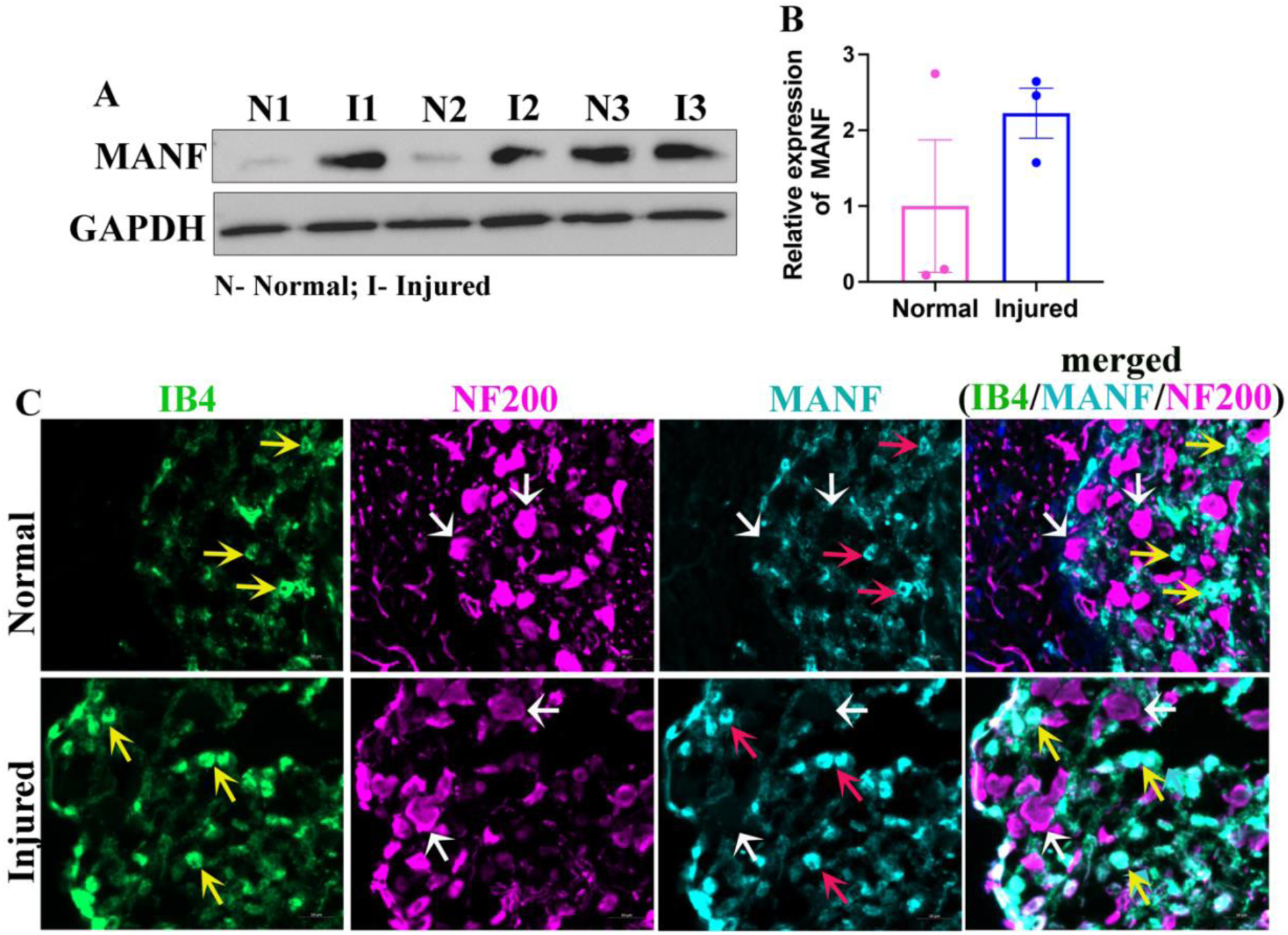
MANF is preferentially expressed in non-peptidergic sensory neurons in adult DRGs. (A) Western blot shows the expression of MANF in the normal and injured (3-day transection injury) adult rat DRGs. GAPDH is used as the loading control. (B) Quantification of ‘A’ shows the induction of MANF in injured DRGs (data presented as mean ± SE; standard ‘t’ test; n=3). (C) Co-immunostaining of IB4 (yellow arrows), NF200 (white arrows) and MANF (red arrows) in normal and injured DRGs shows the preferential expression of MANF in IB4 positive non-peptidergic sensory neurons (yellow arrows in the merged view) (scale bar, 50µm).

### MANF expression is not altered in injured peripheral nerve

We next examined the expression of MANF in injured sciatic nerves to evaluate if MANF is upregulated in nerves after injury. Western blot experiments in non-injured control and injured proximal and distal sciatic nerve segments showed no significant difference in MANF expression, indicating no local induction of MANF in injured nerves (**Figure 2A, B**). Although injury did not induce MANF, a low basal expression of MANF was apparent in both control and injured nerves. Therefore, we next examined the cellular distribution of MANF in nerves by co-immunostaining MANF with βIII tubulin, a pan-neuronal marker, NF200, a large calibre neuron marker, and GFAP, a SC marker. We found that MANF co-localizes with βIII tubulin^+^ axons (**Figure 2C**). βIII tubulin marks both small and large calibre axons. However, we found no notable co-localization of MANF with NF200^+^ axons, which comprise large calibre axons, indicating that MANF is preferentially expressed in NF200^poor^ small calibre axons (**Figure 2D**). Our immunostaining results also showed the co-localization of MANF and GFAP at scattered locations in the nerves indicating that MANF is also expressed occasionally in SCs in both normal and injured peripheral nerves (**Figure 2E)**.

**Figure 2:**
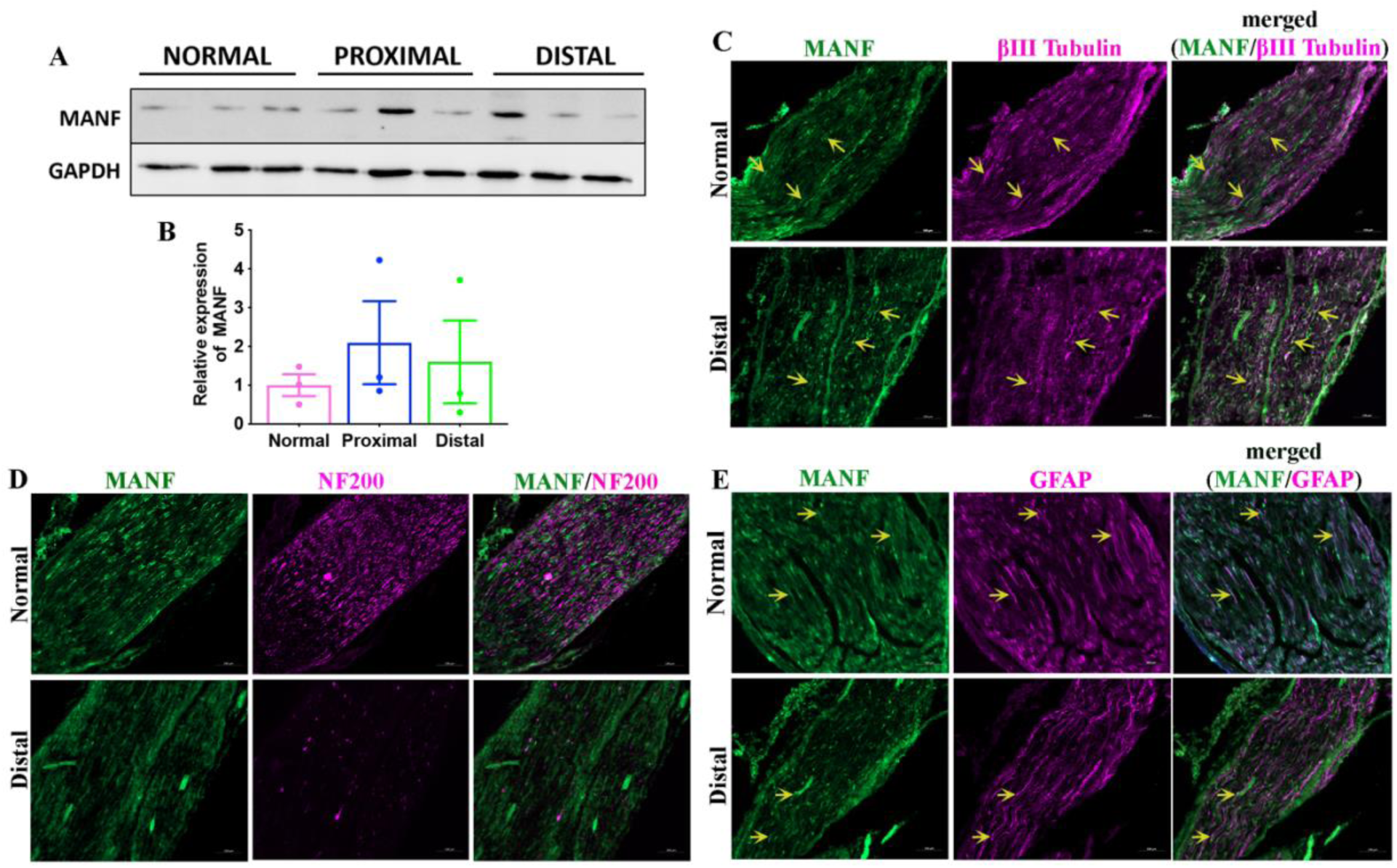
MANF expression does not change in injured peripheral nerves. (A) Western blot shows the expression of MANF in normal and injured (3-day transection injury) proximal and distal sciatic nerve segments of adult rats. GAPDH is used as the loading control. (B) Quantification of ‘A’ shows no significant change in the expression of MANF in injured nerves compared to the control (data presented as mean ± SE; One-Way ANOVA; n=3). (C) Co- immunostaining of βIII tubulin and MANF in normal and injured distal sciatic nerve of adult rats shows MANF expression in a subpopulation of βIII tubulin stained axons (yellow arrows in the merged view) (scale bar, 100µm). (D) Co-immunostaining of NF200 and MANF in normal and injured distal sciatic nerve of adult rats shows no remarkable co-localization of MANF with NF200 positive axons (scale bar, 100µm). (E) Co-immunostaining of GFAP and MANF in normal and injured distal sciatic nerve of adult rats shows expression of MANF in a few populations of SCs (yellow arrows in the merged view) (scale bar, 100µm).

### Exogenous MANF promotes the outgrowth of adult primary sensory neurons *in vitro* regardless of their endogenous MANF expression profile

Our observation of the lack of induction MANF in injured nerves indicates that, although injury slightly induces MANF in DRGs, it is not efficiently transported to injured nerve terminals for axon regeneration. Hence, local supplementation of MANF to injured nerves may be essential for promoting nerve regeneration. Substantiating this argument, we previously showed that exogenous MANF promotes the outgrowth of normal NF200^+^ adult primary sensory neurons *in vitro*(*5*). Here, we investigated if exogenous MANF is even effective in promoting the outgrowth of injured neurons to evaluate its translational potential as a therapy for nerve injury. We also examined if exogenous MANF promotes the outgrowth of all subtypes of neurons by analyzing neurite outgrowth after βIII tubulin staining, which stains both small (NF200^poor^/MANF^high^) and large (NF200^high^/MANF^low^) calibre neurons. We supplemented 50 and 100 ng/ml recombinant human MANF to the cultures of injured and normal adult primary sensory neurons, respectively, as injured neurons have a slightly higher endogenous expression of MANF, as evident from our earlier observation (**Figure 1A, B**). We found that exogenous MANF significantly promoted the outgrowth of both normal and injured primary sensory neurons regardless of the neuron subtypes, as both βIII tubulin^+^ and NF200^+^ neurons showed increased outgrowth after MANF supplementation (**Figure 3A-D**). Overall, these findings substantiate that MANF is a potential therapeutic candidate for peripheral nerve regeneration, requiring systematic therapeutic profiling.

**Figure 3:**
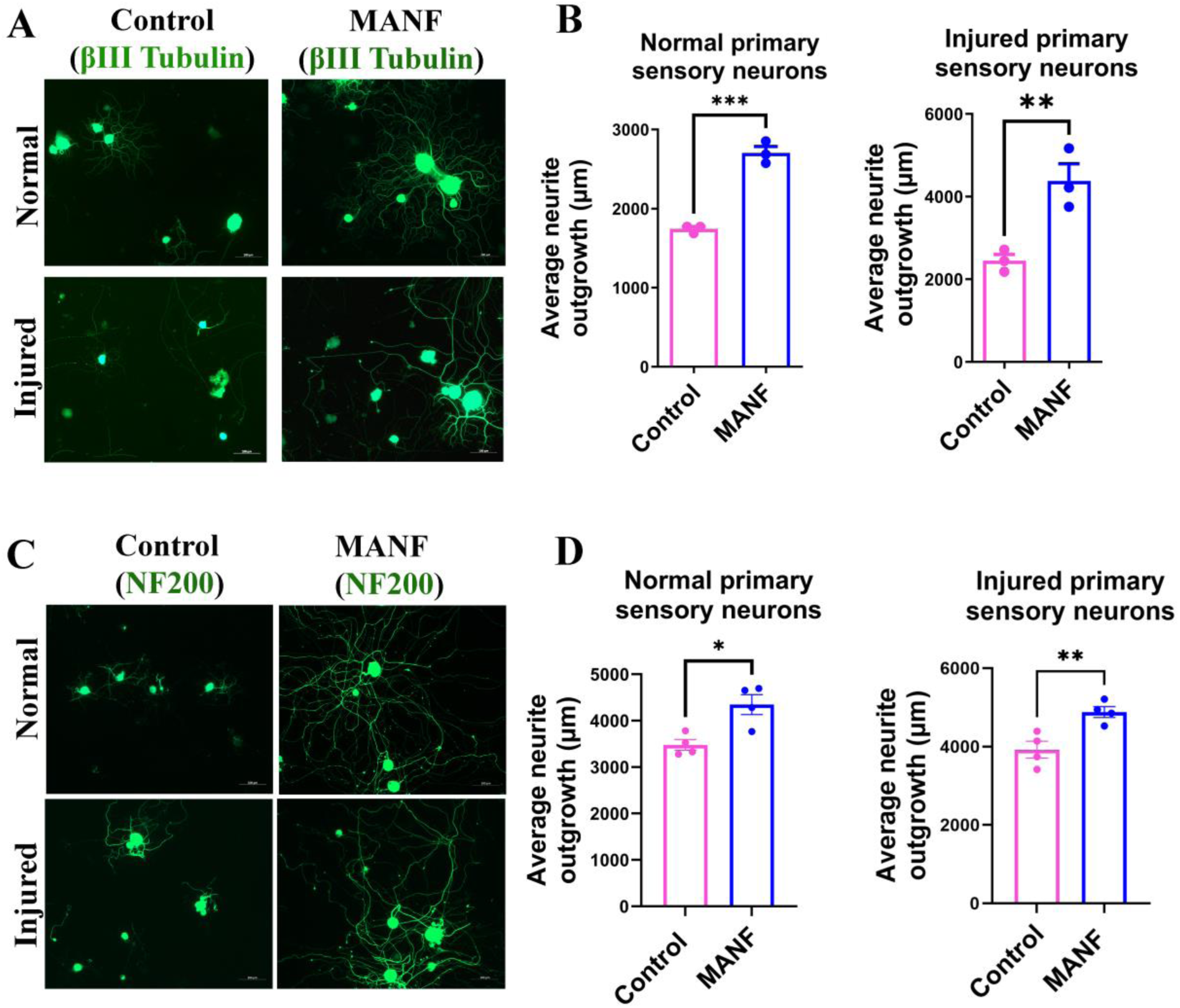
Exogenous MANF promotes the outgrowth of both normal and injured adult neurons *in vitro*. (A) βIII tubulin staining shows that the supplementation of 50 and 100 ng/ml MANF promotes neurite outgrowth of injured and normal adult primary sensory neurons, respectively (scale bar, 100µm). (B) Quantification of neurite outgrowth using WIS-NeuroMath Software shows significant induction of neurite outgrowth in both normal and injured βIII tubulin positive neurons after MANF supplementation (data presented as mean ± SE; Standard ‘t’ test; n=3; **p<0.01, ***p<0.001). (C) NF200 staining shows that the supplementation of 50 and 100 ng/ml MANF promotes neurite outgrowth of injured and normal adult primary sensory neurons, respectively (scale bar, 100µm). (D) Quantification of neurite outgrowth using WIS-NeuroMath Software shows significant induction of neurite outgrowth in both normal and injured NF200 positive neurons after MANF supplementation (data presented as mean ± SE; Standard ‘t’ test; n=4; *p<0.05, **p<0.01).

### Exogenous MANF promotes SC proliferation and migration

SCs are glial cells of the peripheral nervous system. They provide structural and trophic support to regenerating axons. Immediately after a peripheral nerve injury, SCs in the distal nerve stump de-differentiate, proliferate, and then migrate towards the proximal nerve stump and guide the regenerating proximal axons toward their target tissues. These properties of SCs are collectively referred to as ‘SC dynamics’. Therapies designed to improve peripheral nerve regeneration should promote both axon regeneration and SC dynamics. Therefore, we examined the effect of MANF on primary SCs isolated from adult rat sciatic nerves. The purity of the isolated primary SCs was confirmed using GFAP staining (**Figure 4A**). We found a heterogeneous expression profile for MANF in these SCs, with some cells expressing MANF while others showing very low to null expression, which matches our previous observation indicating a low and occasional expression profile for MANF in sciatic nerve resident SCs (**Figure 4B**; **Figure 2E**). MTT assay showed that MANF promotes the proliferation of primary SCs in a dose-dependent manner, with 100 ng/ml MANF inducing a significant increase in proliferation compared to control (**Figure 4C)**. Similarly, the scratch assay showed improved gap closure after MANF supplementation indicating that MANF promotes the migration of primary SCs (**Figure 4D, E)**. Improved gap closure in a scratch assay may also result from increased proliferation of cells. Therefore, we performed a transwell migration assay to confirm the ability of MANF to promote SC migration (**Figure 4F)**. As expected, we found increased migration of SC across the membrane, towards the MANF compartment, in the transwell assay, confirming the ability of MANF to promote SC migration by providing tropic signaling (**Figure 4G,H)**.

**Figure 4:**
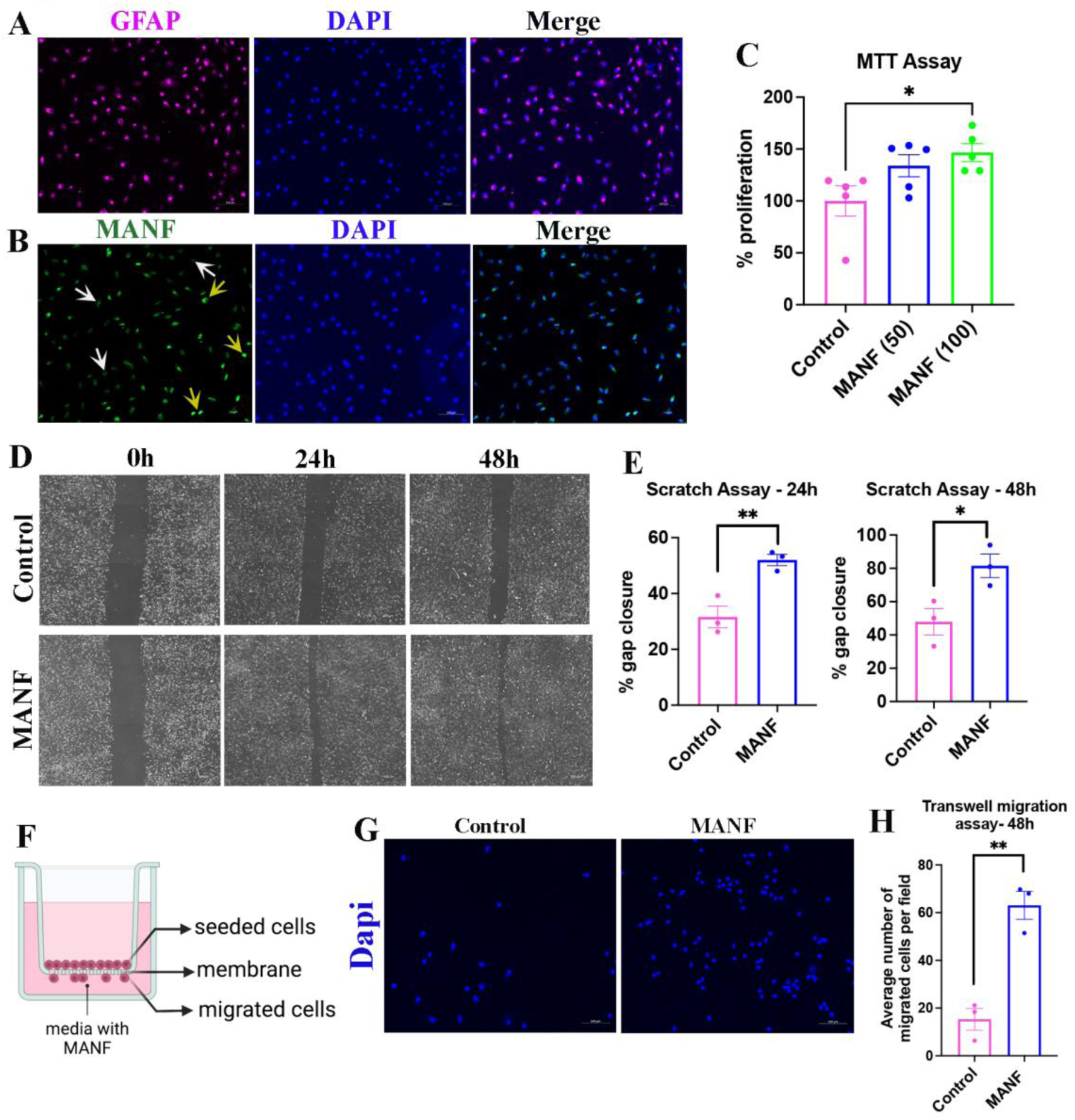
Exogenous MANF promotes SC dynamics. (A) Immunostaining of GFAP in adult rat primary SCs (scale bar, 100µm). (B) Immunostaining of MANF in adult rat primary SCs. Yellow arrows show intense expression, and white arrows show weak to null expression (scale bar, 100µm). (C) MTT assay using adult rat primary SCs shows increased proliferation of cells in response to exogenous MANF supplementation (data presented as mean ± SE; One-Way ANOVA (Tukey’s multiple comparisons test); n=6; *p<0.05). (D) Brightfield images of the scratch assay using adult rat primary SCs show a time-dependent gap closure in control and MANF (50 ng/ml) supplemented groups (scale bar, 100µm). (E) Quantification of percentage gap closure in scratch assay shows increased gap closure in MANF (50 ng/ml) supplemented group compared to control (data presented as mean ± SE; Standard ‘t’ test; n=3; *p<0.05, **p<0.01). (F) Schematic of transwell migration assay. (G) DAPI staining on the lower side of the membrane from a transwell migration assay shows migrated adult primary SCs (scale bar, 100µm). (H) Quantification of transwell migration assay shows increased migration of primary SCs in MANF (50 ng/ml) supplemented group (data presented as mean ± SE; standard ‘t’ test; n=3; **p<0.01).

Next, we repeated the MTT and transwell migration assays in the S16 SC line and found that supplementation of MANF promotes the proliferation and migration of these cells, too (**Figure S1**). Overall, our result indicates that MANF promotes the dynamics of SCs that are desirable for peripheral nerve regeneration.

### Local and repeated administration of MANF promotes axon regeneration *in vivo*

Potent nerve repair therapies, when seamlessly available to injured nerves, should promote axon regeneration. Therefore, to test if MANF delivered locally to injured nerves promotes axon regeneration, we directly injected MANF into adult CD1 mouse sciatic nerves after a crush injury. Crush injuries represent the most frequent type of nerve injuries presented in neuro-clinics. A crush injury induces distal axon degeneration, with partial or no disruption of the connective tissue layers of nerves. We directly injected three doses of MANF (2µg) at the site of the crush injury, with the first dose given immediately after the crush injury, the second dose on day 3, and the third dose on day 5 (**Figure 5A**). The nerves were harvested on day 7, and the regenerating axons in the distal nerve were quantified after βIII tubulin staining (**Figure 5B**). We found an increased number of intact axons in the distal nerve in the MANF group compared to the control (saline), indicating that locally administered MANF promotes axon regeneration *in vivo* (**Figure 5C, D**). In contrast, the control group had more degenerating axon profiles observed in the distal nerve compared to the MANF group, suggesting that clearing of the degenerated axonal debris is also faster in the MANF group.

**Figure 5:**
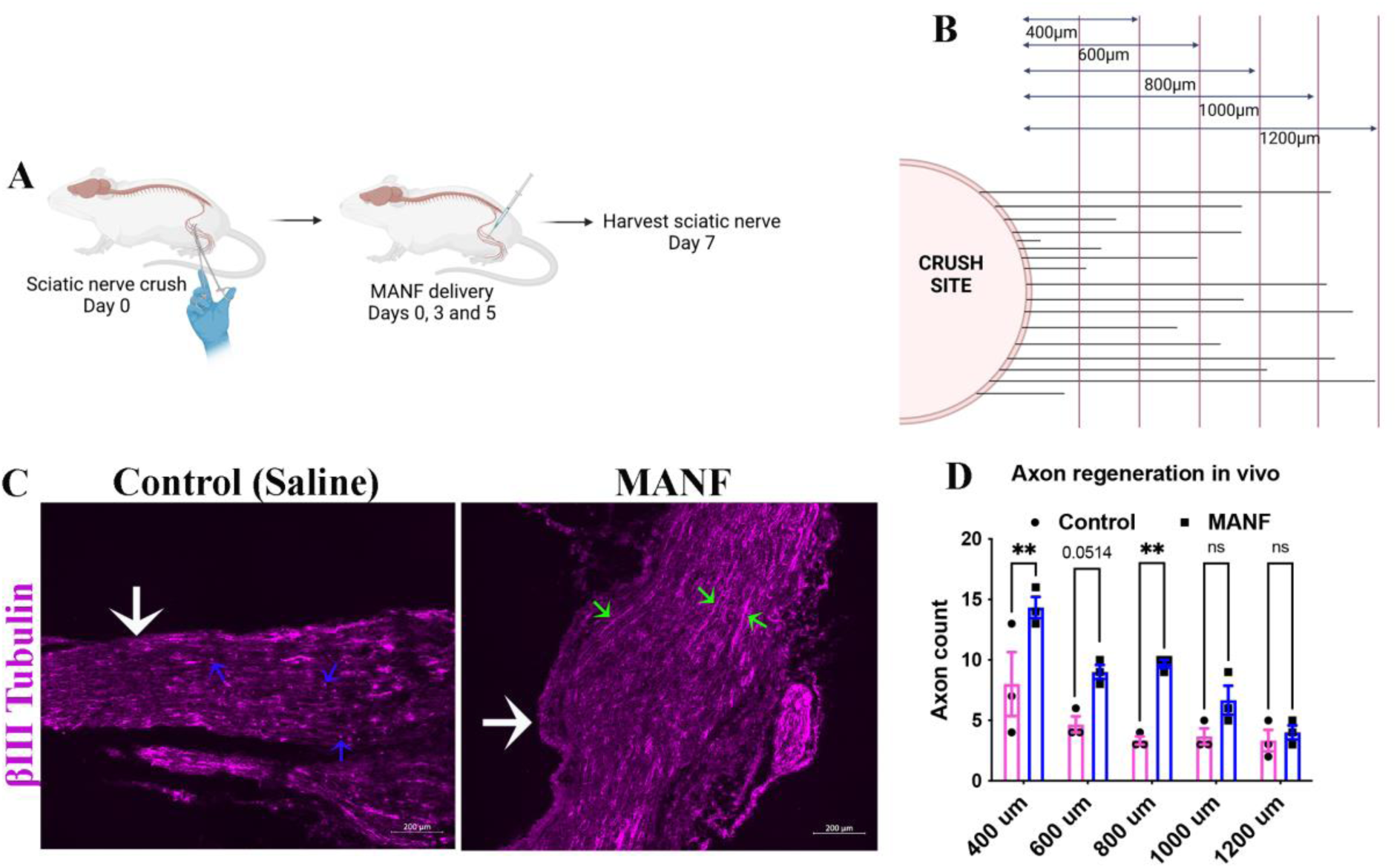
Local and multiple supplementations of MANF to injured nerve promote axon regeneration. (A) Schematic of *in vivo* nerve crush injury and local supplementation of MANF in adult mice. (B) Schematic of axon quantification approach. The number of axons crossing the perpendicular lines drawn at the shown distances are manually counted to tabulate the axon count at each distance. (C) βIII tubulin staining of sections of injured sciatic nerves from adult mice shows degenerated (blue arrows) and intact (green arrows) axons in the control and MANF group, respectively. White arrows show the crush site (scale bar, 200µm). (D) Quantification of intact axons in the distal nerve segment shows an increased number of axons in the MANF group compared to the control at varying distances from the crush site (data presented as mean ± SE; Two-Way ANOVA (Sidak’s multiple comparisons test); n=3; **p<0.01).

### Nerve resident SCs expressing Doxycycline (Dox)-inducible MANF protect axons and promote regeneration

Our findings shown above suggest that local and frequent supplementations of MANF to nerves promote axon regeneration *in vivo*. However, surgical re-exposure of nerves for repeated delivery of MANF, as done for the proof-of-principle experiment shown in **Figure 5**, is challenging in clinical settings. Therefore, we examined if nerve resident SCs can be exploited as a local and sustained delivery system for MANF. To explore this possibility, we first expressed Dox-inducible MANF in cultured primary SCs using lentiviral transductions (**Figure S2**). We found that the generated SC-MANF grows healthy on regular culture surfaces and an artificial nerve conduit (**Figure 6A, B)**. ELISA assay confirmed that SC-MANF secrete significantly higher levels of MANF as compared to lenti-empty vector (L-EV) transduced SCs following Dox exposure (**Figure 6C)**.

**Figure 6:**
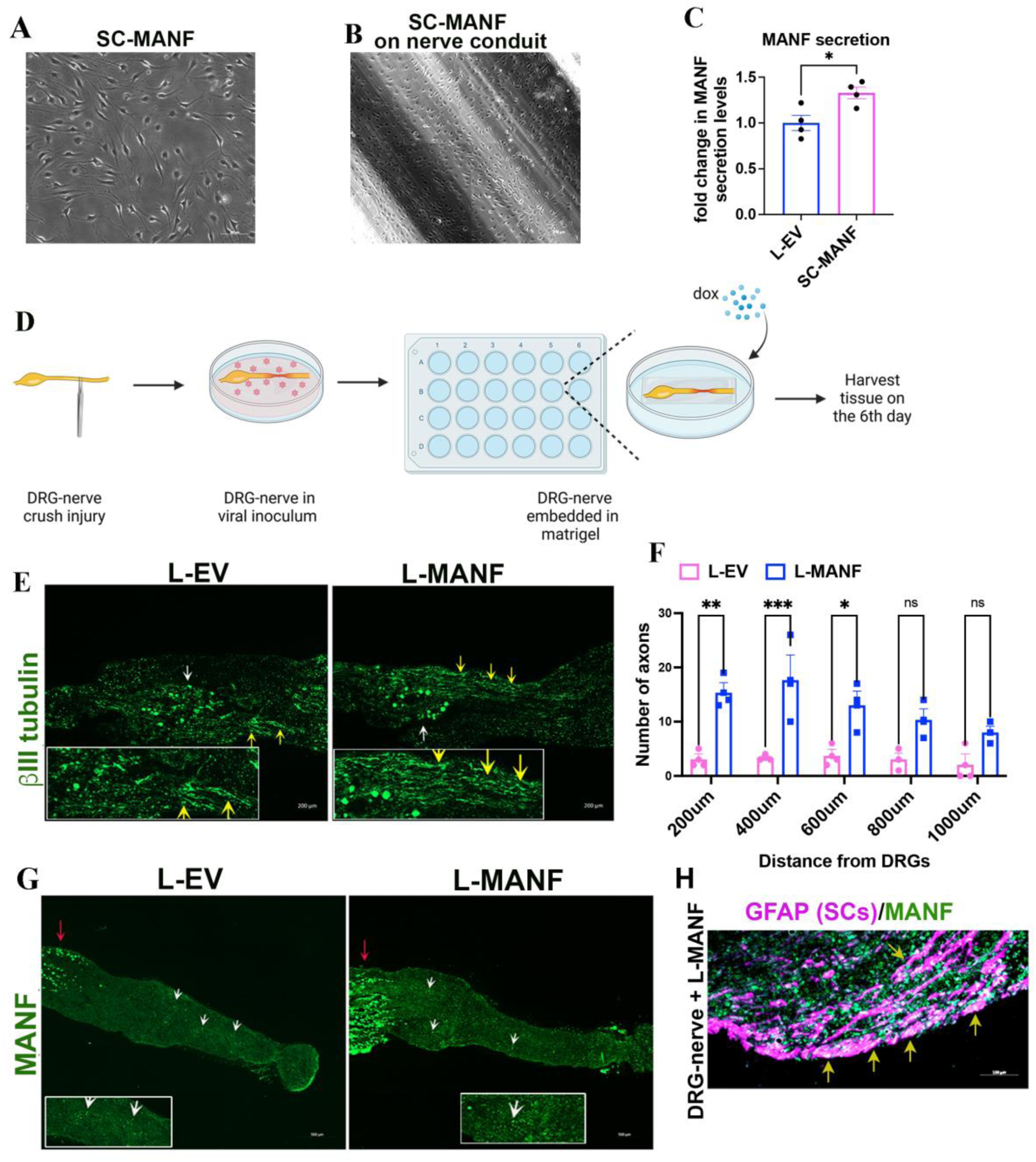
SC expressing Dox-inducible MANF (SC-MANF) protects axons and promotes regeneration. (A) SC-MANF after seven days of puromycin selection in a culture flask. **(B**) SC- MANF grows healthy on an artificial nerve conduit (15-day). (C) ELISA assay shows increased secretion of MANF in the SC-MANF group compared to L-EV transduced primary SCs after Dox supplementation (data presented as mean ± SE; standard ‘t’ test; n=4). (D) Schematic of nerve crush injury experiment using DRG-nerve explant. (E) Immunostaining of βIII tubulin in sections of crush injured DRG-nerve explant (day 6) showing intact axons (yellow arrows) in the L-EV and L-MANF transduced groups. The white arrow shows DRG (scale bar, 200µm). (F) Quantification of axons in DRG-nerve explants on day 6 shows an increased number of intact axons in the L- MANF group compared to the L-EV group (data presented as mean ± SE; Two-Way ANOVA (Sidak’s multiple comparisons test); n=3; *p<0.05, **p<0.01, ***p<0.001). (G) Immunostaining shows the expression of MANF in L-EV and L-MANF transduced DRG-nerve explants cultured in the presence of Dox (1µg/ml). White arrows show the expression of MANF (green dots) in the nerve segment, and the red arrow shows MANF expression in DRG (scale bar, 500µm). Enlarged areas are provided in the insets. (H) A representative image shows the expression of MANF in SCs (SC-MANF generation) in the nerve segment of a DRG-nerve explant transduced with L-MANF and cultured in the presence of Dox.

Next, to examine if nerve-resident SCs expressing Dox-inducible MANF (SC-MANF) promote axon regeneration, we used DRG-nerve explants isolated from adult SD rats. A nerve crush injury was made to the explants, and they were then transduced with either L-EV or L- MANF (to generated nerve-resident SC-MANF) and cultured in Cultrex extracellular membrane for six days in the presence of Dox (**Figure 6D)**. As opposed to nerve crushes *in vivo*, a nerve crush to DRG-nerve explant induces degeneration of proximal axons, especially during the first week of injury. Interestingly, the L-MANF group showed an increased number of intact proximal axons on day 6 compared to the L-EV group (**Figure 6E, F**). The greater number of intact axons in the L- MANF group may result from the combined neuroprotective and regenerative actions of MANF secreted by the generated, resident SC-MANF in the explants. Immunostaining confirmed the increased induction of MANF in the L-MANF group compared to L-EV group after Dox exposure (**Figure 6G**). We also confirmed the integration of L-MANF in nerve resident cells, including SCs *(generation of nerve-resident SC-MANF)*, throughout the nerve (**Figure 6H**)

We next repeated the explant experiment for 15 days for selectively capturing regenerating axons crossing the crush site (**Figure 7A**). GAP-43 staining showed the presence of significantly increased number of regenerating axons in the distal nerves in the L-MANF group compared to the L-EV group, indicating significant nerve regeneration **(Figure 7B).** We found even distribution of SCs in distal nerves in the L-MANF group, indicating that SC integrity is maintained well in this group **(Figure 7C).** As observed in 6-day cultures, immunostaining of 15-day cultures showed an increased expression of MANF throughout the nerve in the L-MANF group compared to L-EV group after Dox exposure, confirming the induction of MANF in the L-MANF group (**Figure 7D**). Collectively, these results show that nerve-resident SC-MANF is a potential cellular therapy for peripheral nerve regeneration.

**Figure 7:**
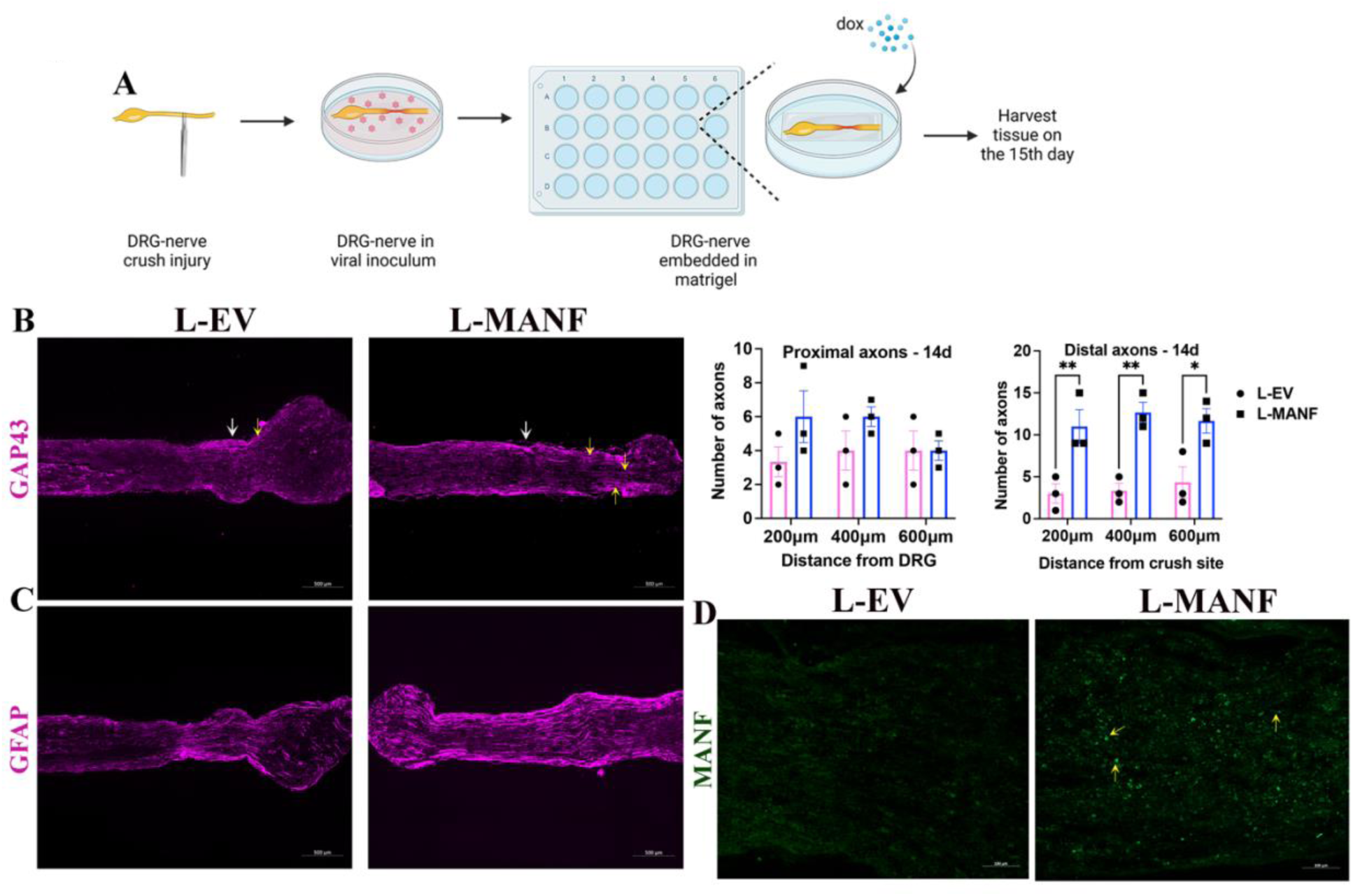
SC-MANF promotes axon regeneration. (A) Schematic and timeline of the nerve crush injury experiment using DRG-nerve explants. (E) Immunostaining of GAP-43 in sections of DRG-nerve explants (day 15) shows longer regenerating axons (yellow arrows) past crush site (white arrow) in the L-MANF group compared the L-EV group (scale bar, 500µm). (B) Quantification of regenerating proximal and distal axons in DRG-nerve explants on day 15 shows a significant increase in regenerating distal axons in the L-MANF group compared to L-EV group (data presented as mean ± SE; Two-Way ANOVA (Sidak’s multiple comparisons test); n=3; *p<0.05, **p<0.01). (C) GFAP staining in sections of DRG-nerve explants (day 15) shows healthy distribution of SCs in the distal nerve of L-MANF group (scale bar, 500µm). (D) Immunostaining shows the expression of MANF in L-EV and L-MANF transduced DRG-nerve explants cultured in the presence of Dox (1µg/ml) for 15 days. Yellow arrows show the expression of MANF (green dots) in the nerve segment (scale bar, 100µm).

## DISCUSSION

Successful regeneration of peripheral nerves requires the coordinated actions of axons and SCs(3, 10). Although such coordinated activities exist for a few days to weeks following a nerve injury, the axons and SCs gradually lose mutual growth support, especially in the distal nerve segment, due to the poorly regrowing axons failing to re-establish contact with SCs. Therapies that can improve the pace of axon growth and sustain the growth competence and dynamics of axons and SCs, respectively, should be able to maintain the axon-SCs mutual interaction and facilitate regeneration. In this work, we demonstrated that MANF is a potential therapy that can promote both axon growth and SC dynamics to sustain regeneration.

MANF is a neurotrophic factor(11, 12). Previous research showed that neurotrophic factors, such as nerve growth factor (NGF), can promote regeneration in animal models(13). However, translating NGF into a clinical therapy posed challenges because of its unexpected side effects and difficulties in maintaining its therapeutic levels locally in injured nerves (2). NGF was also shown to induce pain, making it a less favorable choice for clinical therapy (14). Although MANF is a neurotrophic factor, it is structurally and functionally distinct from the NGF family of neurotrophic factors(15). Also, NGF uses tropomyosin kinase receptor A (TrkA) for its growth promoting actions, while a specific receptor for MANF’s neurotrophic actions is unknown. This suggests the involvement of an alternate growth promoting mechanism for MANF, independent of TrkA, raising its translational potential as a clinical therapy.

While neurotrophic factors are attractive targets for nerve regeneration, their supraphysiological concentrations may suppress axon growth(16). Hence, sustained but optimal delivery of neurotrophic factors to injured nerves is critical for maintaining the momentum of axon growth. We initially examined the effect of intraperitoneal administration of MANF on nerve regeneration in animal models of sciatic nerve crush injury but did not find any significant improvement in nerve regeneration and functional recovery (data not presented). However, local and repeated administration of MANF to injured nerves for a short term has been shown to improve axon growth after a crush injury. This is promising and detailed studies covering longer time points and functional recovery assessments are warranted to demonstrate the full potential of MANF as a nerve repair therapy.

In addition to its neurotrophic actions, MANF modulates cellular stress by combating unfolded protein response (UPR) (17, 18). This UPR buffering effect of MANF is linked to its neuroprotective effect (19). For example, experimental models of Alzheimer’s Disease, Parkinson’s Disease and stroke have demonstrated that MANF is a neuroprotector (20–22). While our *in vitro* neurite outgrowth experiments showed that MANF promotes regeneration, the effect of MANF in maintaining intact axons in our explant model may be attributed to its combined neuroprotective and regenerative actions. Notably, our explant model shows proximal axon degeneration in the first week period, while such degenerative events were comparatively lesser in the MANF group compared to the control, indicating neuroprotection. At the same time, the 15- day axon regeneration study time point in the explant model showed longer distal regenerating axons in the L-MANF group compared to the control demonstrating the potential of MANF to improve nerve regeneration. Similarly, MANF promoted axon growth past proximal stump in our *in vivo* crush injury model also, further substantiating the neuroregenerative properties of MANF.

Our *in vivo* experiment was primarily focused on the effect of local MANF on nerve regeneration and we found that local and repeated supplementation of recombinant MANF in nerves improves axon regeneration. A previous study by another group demonstrated that decellularized human nerve grafts seeded with MANF expressing mesenchymal stem cells improve nerve regeneration and functional recovery in adult rates (6). While we and this group used different nerve injury models *(crush injury vs transection injury)* and MANF delivery approaches *(direct administration of MANF vs allogenic transplantation of mesenchymal stem cells expressing MANF),* both these studies showed that locally available MANF improves axon regeneration, substantiating the potential of MANF as a neurotherapeutic. In this work, we additionally demonstrated that MANF promotes SC dynamics that are favorable for regeneration, and thus, revealed an excellent opportunity to use autologous SCs as a carrier system for the local delivery of MANF. SCs are an integral component of peripheral nerves, and their partnership with axons is critical for the proper functioning of nerves. Here we showed that SCs programmed for stably expressing MANF provide an excellent opportunity to timely and temporally deliver MANF to injured nerves. We also showed that this approach protects axons and promotes regeneration. Importantly, the use of programmed SCs expressing MANF as a cellular therapy is comparatively safer than stem cell therapies, as autologous SCs carry no risk of immune rejection. However, further refinements are required to this approach, such as tailoring the Dox dosage regimen to accomplish the most optimal levels of MANF in regenerating nerves and evaluating this approach in chronic injury models, for the complete profiling of SC-MANF as a local therapy for peripheral nerve regeneration.

Overall, we demonstrate that MANF protects peripheral axons and induces a favorable growth response in both axons and SCs to improve peripheral nerve regeneration. We further showed that local supplementation of MANF to injured nerves at regular intervals promotes regeneration. We designed a potential cellular therapy for nerve regeneration by harnessing the capacity of SCs as a carrier system for the local and sustained delivery of MANF. Further refinement of this potential therapy in chronic nerve injury models will ensure a novel therapeutic opportunity for peripheral nerve regeneration.

## Acknowledgements

This work is supported by the College of Medicine Research Award from the College of Medicine at the University of Saskatchewan to AK (Krishnan). This work is also partly supported by the Natural Sciences and Engineering Research Council (NSERC) of Canada Discovery Launch Supplement (DGECR-2020-00049) to AK. We also acknowledge the CoMGRAD fellowship from the College of Medicine to BS.

## Author contributions

BS generated and analyzed data and contributed to the preparation of the initial and revised versions of the manuscript. CH, VM, and NJ contributed to the data generation and analysis. AK (Kumar) and JVJ provided resources for lentiviral and nerve conduit experiments, analyzed data, and contributed to the preparation of the initial and revised versions of the manuscript. AK (Krishnan) conceived the concept, generated the resources for the research, analyzed the data, and contributed to the preparation of the initial and revised versions of the manuscript and finalized the manuscript.

## Supplementary figure legends

**Figure S1:**
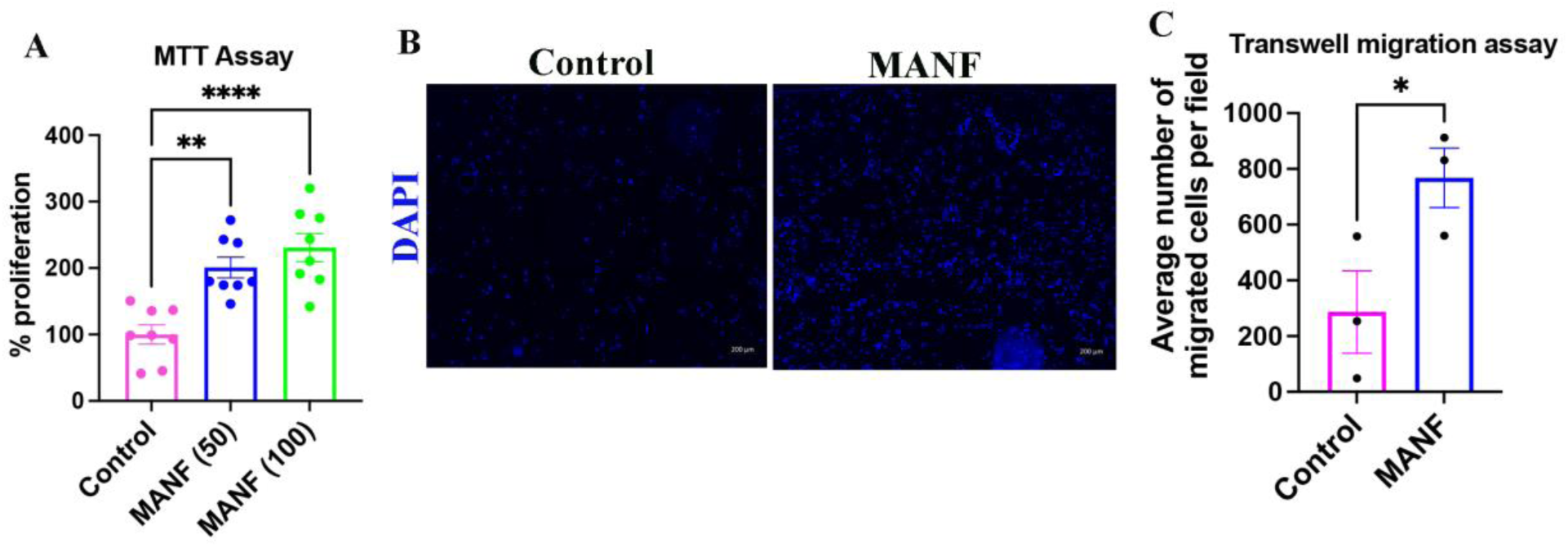
Exogenous MANF promotes SC dynamics. (A) MTT assay using S16 SC line shows increased proliferation of cells in response to exogenous MANF (data presented as mean ± SE; One-Way ANOVA (Tukey’s multiple comparisons test); n=8; **p<0.01, ****p<0.0001). (B) DAPI staining on the lower side of the membrane from a transwell migration assay shows migrated S16 SCs at 48h in the control and MANF group (scale bar, 200µm). (C) Quantification of transwell migration assay shows increased migration of S16 SCs in MANF (100 ng/ml) supplemented group at 48h (data presented as mean ± SE; standard ‘t’ test; n=3; *p<0.05).

**Figure S2:**
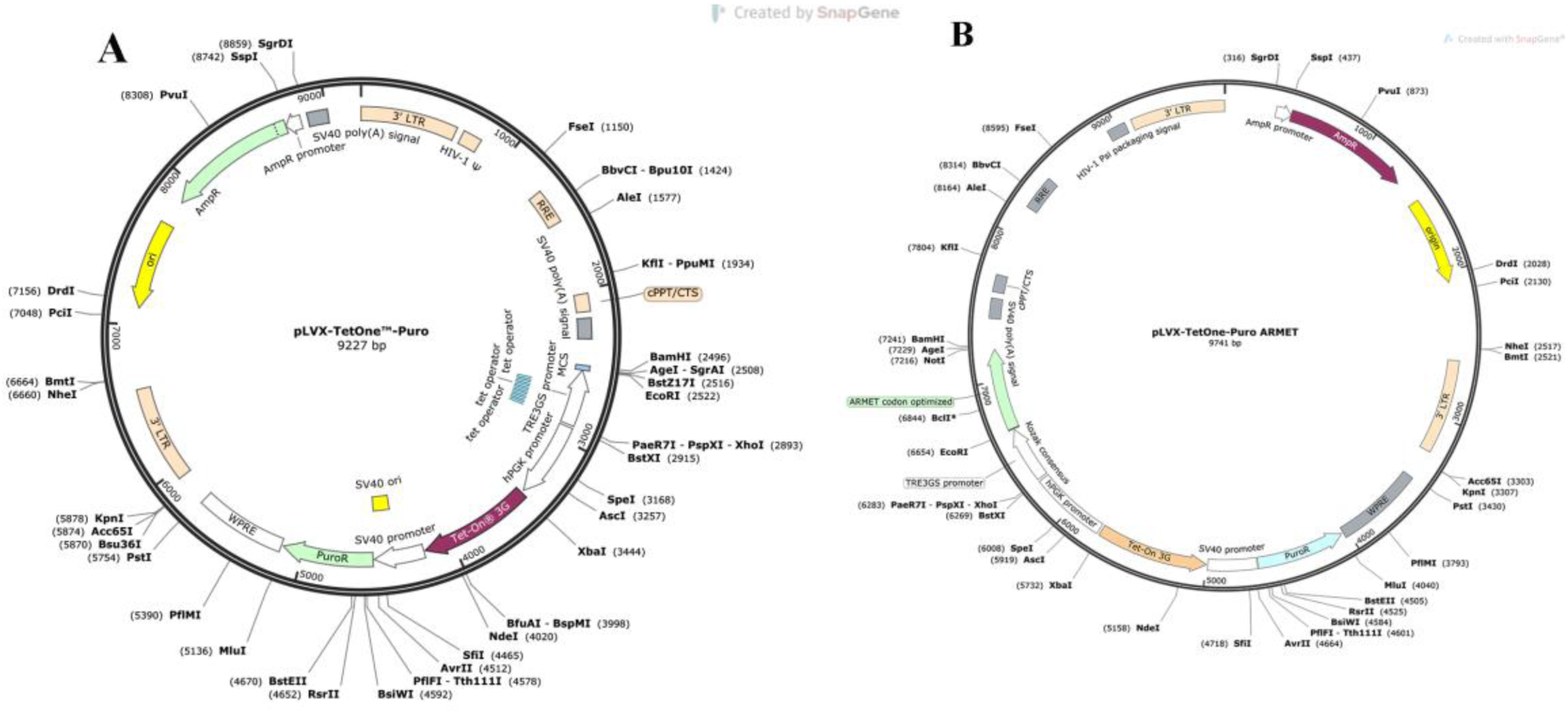
Maps of (A) pLVX TetOne Puro (empty vector) and (B) pLVX TetOne Puro MANF constructs.

